## Supplementary Figures for "Structure and activation mechanism of the Makes caterpillars floppy 1 toxin"

### Supplementary information

**Supplementary Movie 1. Architecture of Mcf1.**

**Supplementary Movie 2. Mechanism of activation of Mcf1.**

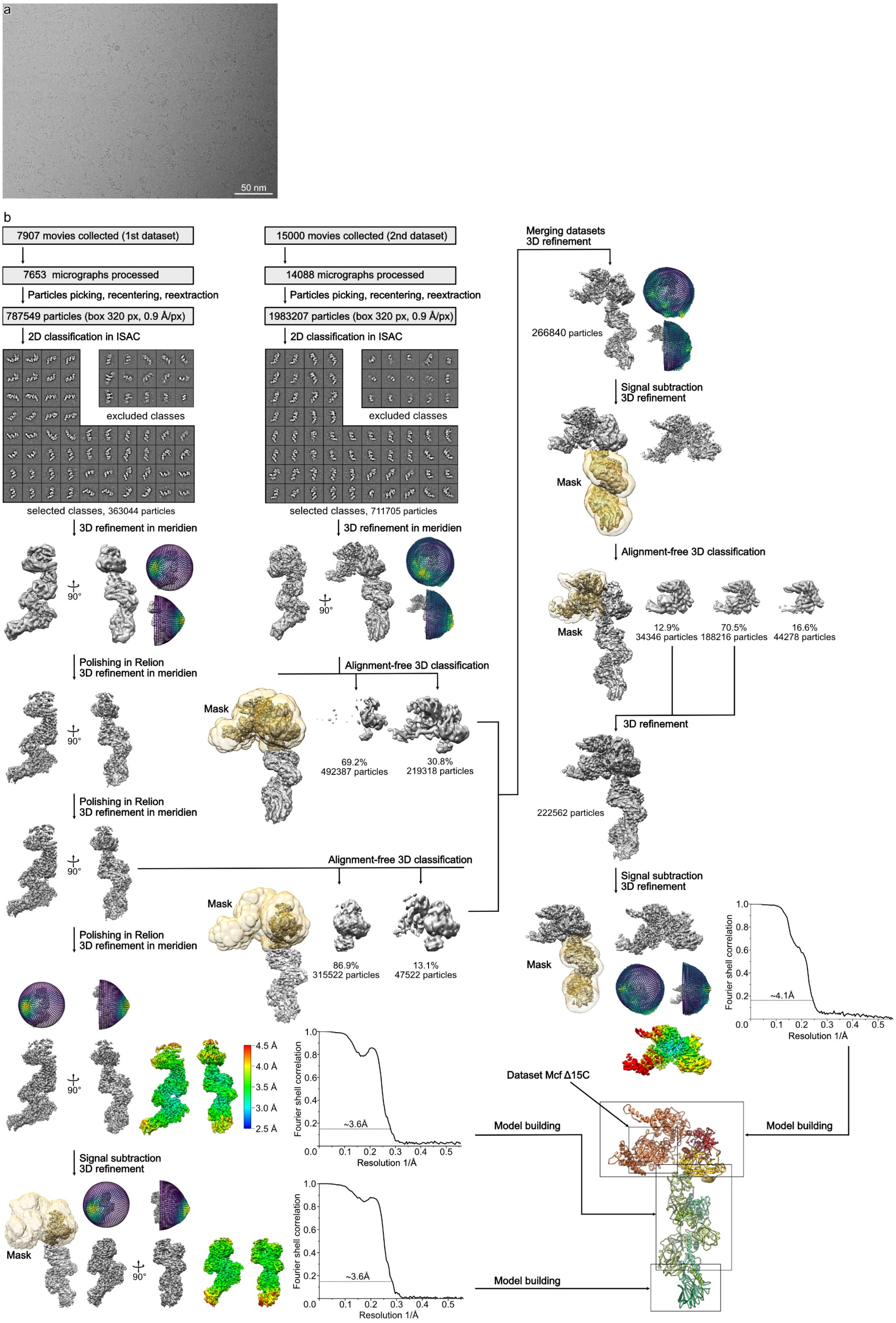

**Supplementary Fig. 1.** Processing of the full-length Mcf1 dataset. **a** An example of analyzed cryo-EM micrographs. **b** Processing overview with angular distributions and Fourier shell correlation curves of the final reconstructions used for model building.

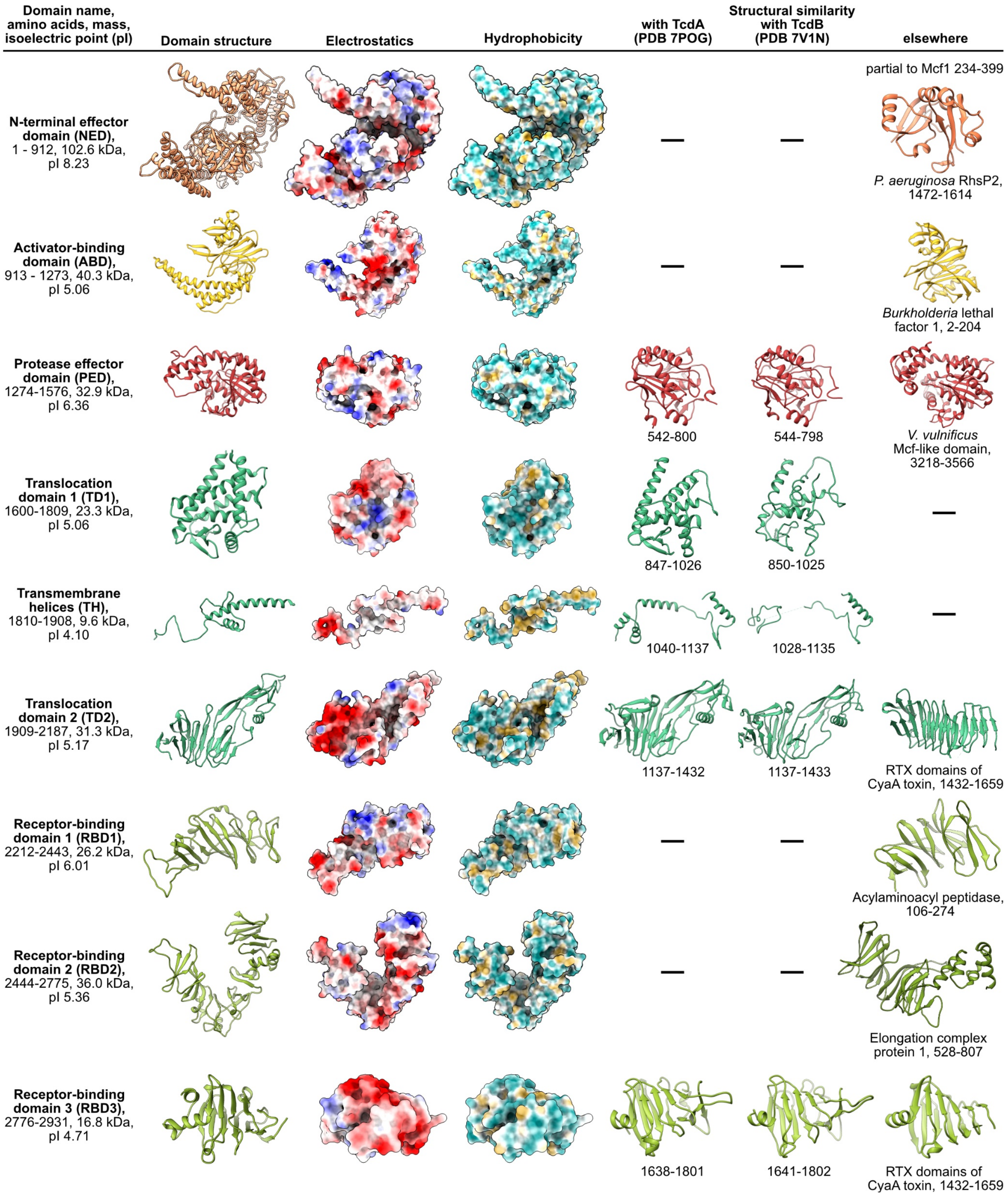

**Supplementary Fig. 2.** Domains of the Mcf1 toxin and their structural homologues (*C. difficile* TcdA toxin – PDB 7POG[1]; *C. difficile* TcdB toxin – PDB 7V1N[2]; *P. aeruginosa* RhsP2 toxin – PDB 7RT7[3]; *Burkholderia* lethal factor – PDB 3TU8[4]; *Vibrio vulnificus* Mcf-like domain – PDB 6II6[5]; RTX domains of *Bordetella pertussis* CyaA toxin – PDB 6SUS[6]; Acylaminoacyl peptidase – PDB 3O4J[7]; Elongation protein complex 1 – PDB 6QK7[8]).

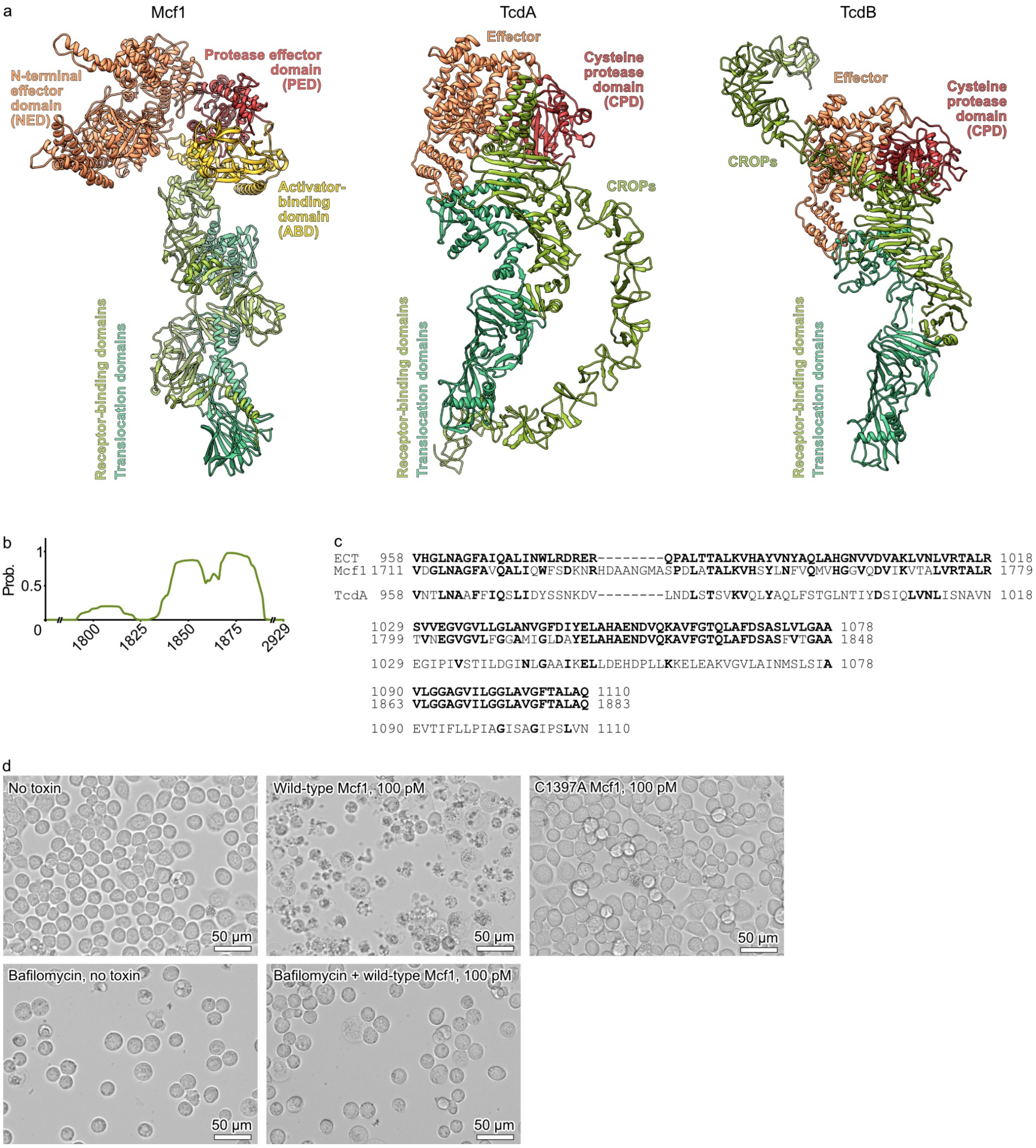

**Supplementary Fig. 3.** Mcf1 is a member of the superfamily of large clostridial toxins. **a** Structural comparison of Mcf1 with TcdA (PDB 7POG[1]) and TcdB (PDB 7V1N[2]). **b** Computational analysis of potential transmembrane regions in Mcf1 by TMHMM 2.0 [9]. Prob. – amino acid probability score of being part of a transmembrane helix. **c** Consensus sequence alignment of evolutionary conserved translocase (ECT) of large clostridial toxins, Mcf1 and TcdA. **d** Toxicity of Mcf1 in insect Sf9 cells. The experiment was performed twice.

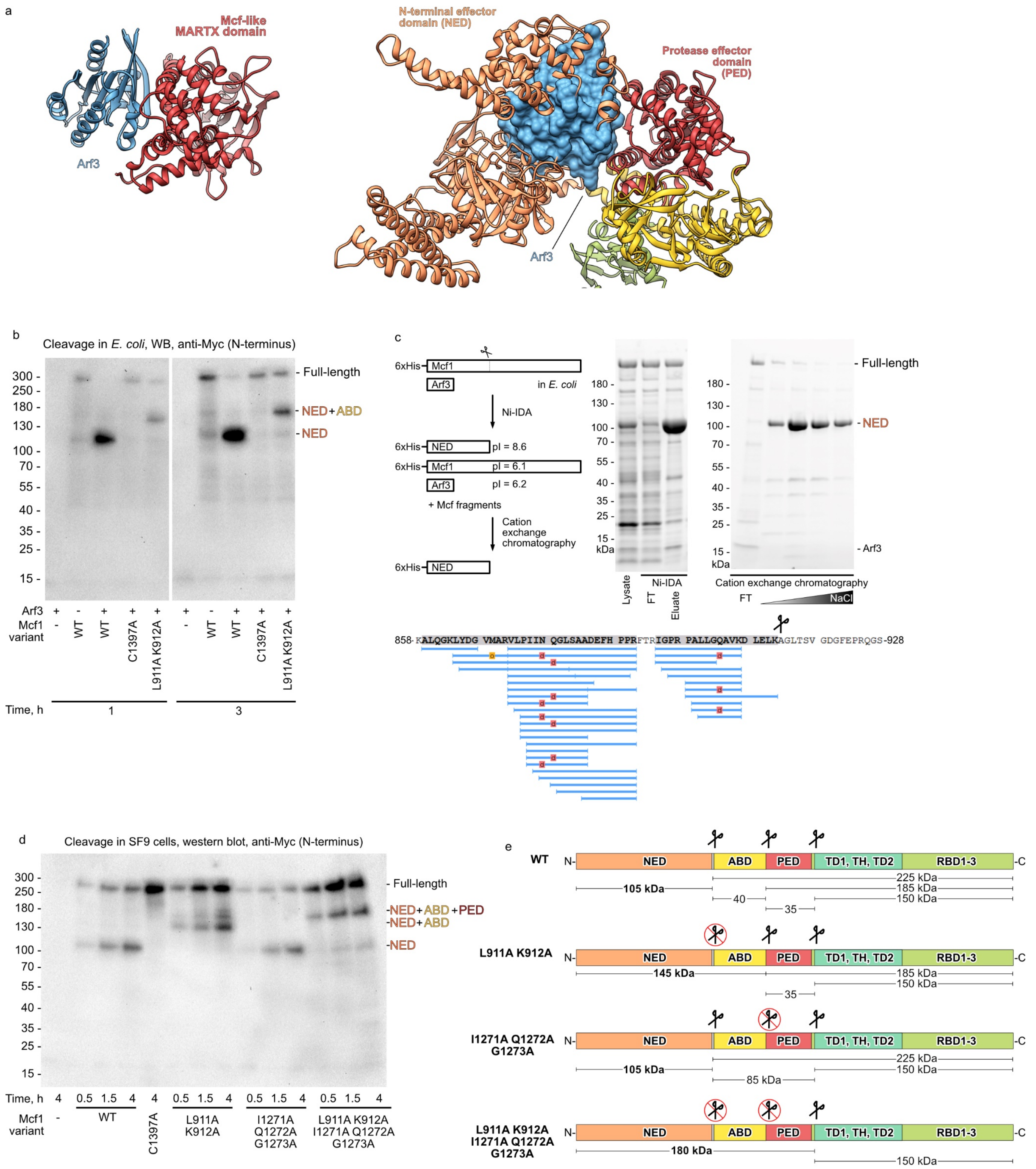

**Supplementary Fig. 4.** Autoprocessing of Mcf1. **a** Arf3 cannot bind to the protease effector domain of Mcf1 in the same way that it binds to the Mcf-like MARTX domain (PDB 6II6, [5]) due to the sterical clash with the N-terminal effector domain. **b** Western blot (WB) analysis of lysates of *E. coli* coexpressing Arf3 and Myc-tagged Mcf1 variants. **c** Purification of the N-terminal effector domain (NED) after coexpression of Arf3 and the full-length Mcf1 in *E. coli* (above) and the peptide coverage of the C-terminal region of the effector in liquid chromatography with tandem mass spectrometry (below). **d** Western blot analysis of N-terminally Myc-tagged Mcf1 autoproteolysis during cell intoxication. All SDS-PAGE and western blot analyses were performed twice. **e** Schematic illustration of the Mcf1 cleavage pattern. IDA - Iminodiacetic acid. Abbreviations: see Fig. 1.

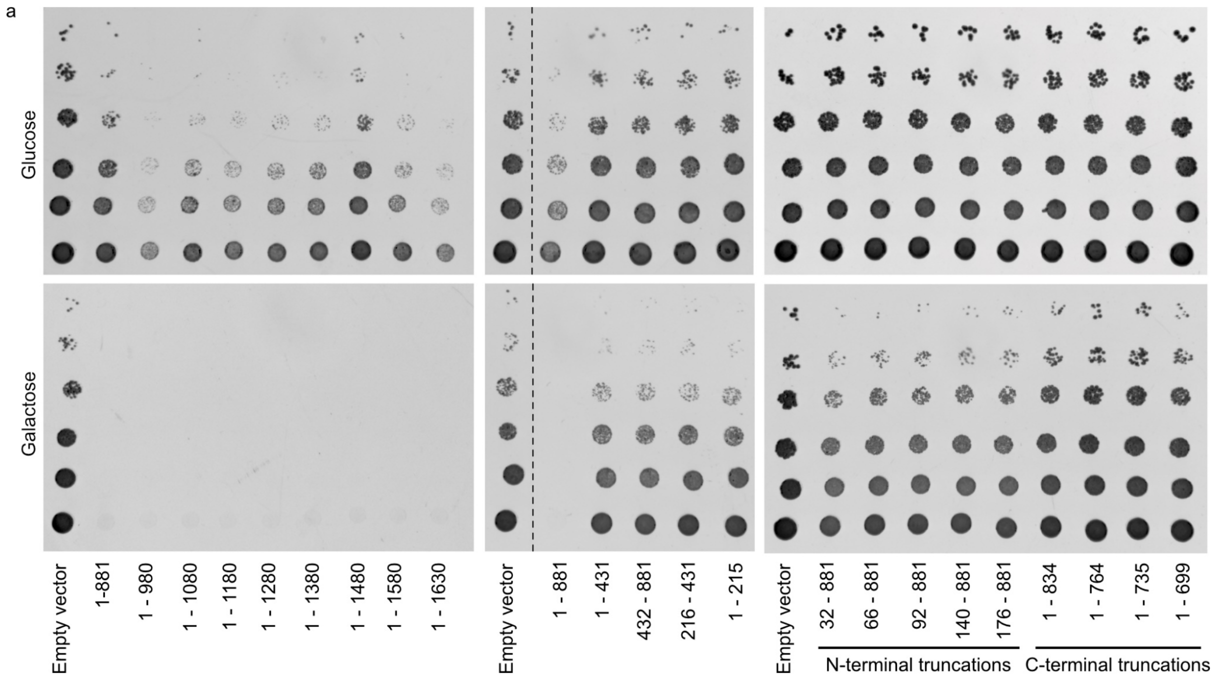

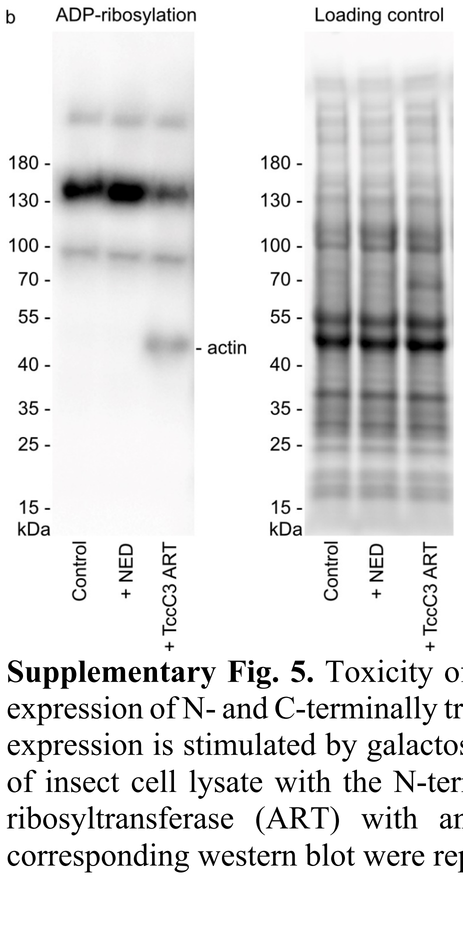

**Supplementary Fig. 5.** Toxicity of Mcf1 fragments in *S. cerevisiae*. **a** Yeast viability upon expression of N- and C-terminally truncated Mcf1 fragments. In yeast drop tests, *mcf1* fragment expression is stimulated by galactose, and repressed by glucose. **b** ADP-ribosylation analysis of insect cell lysate with the N-terminal effector domain (NED) of Mcf1 and TccC3 ADP-ribosyltransferase (ART) with anti-ADP-ribose binding protein. The reaction and the corresponding western blot were repeated twice.

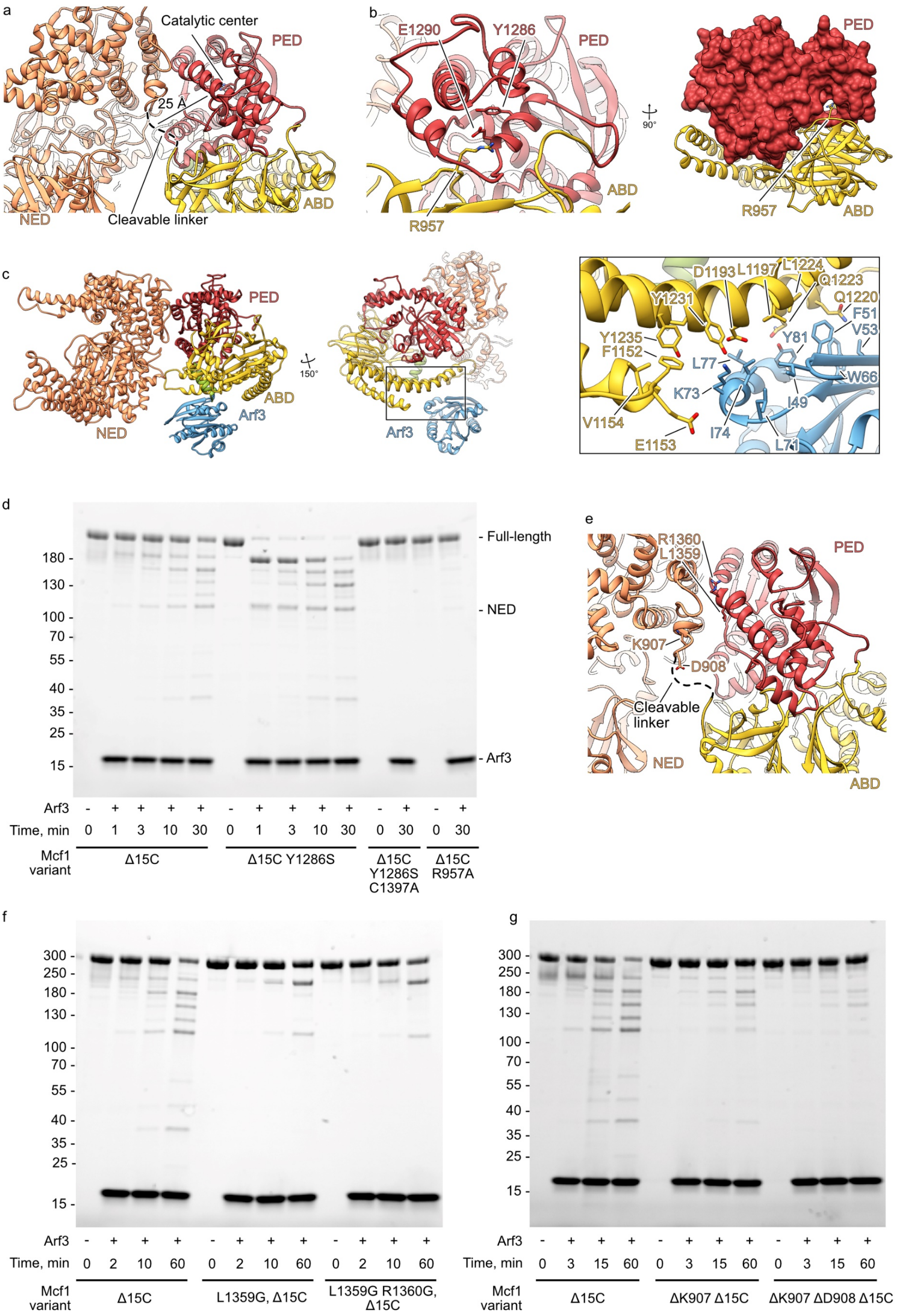

**Supplementary Fig. 6.** Mechanism of Mcf1 activation. **a** The cleavable linker is localized in 25 Å distance from the catalytic center of the protease effector domain (PED). **b** The structure of the hook (R957) – loop hinge of Mcf1. **c** Structure of the Mcf1$_{C1397A\ \Delta15C}$-Arf3 complex with a focus on the interaction interface. **d, f, g** Autoproteolysis of Mcf1 variants in the presence of Arf3. **e** Position of the cleavable linker-preceding amino acids (K907, D908) and amino acids that create a rigid scaffold of the system (L1359, R1360).

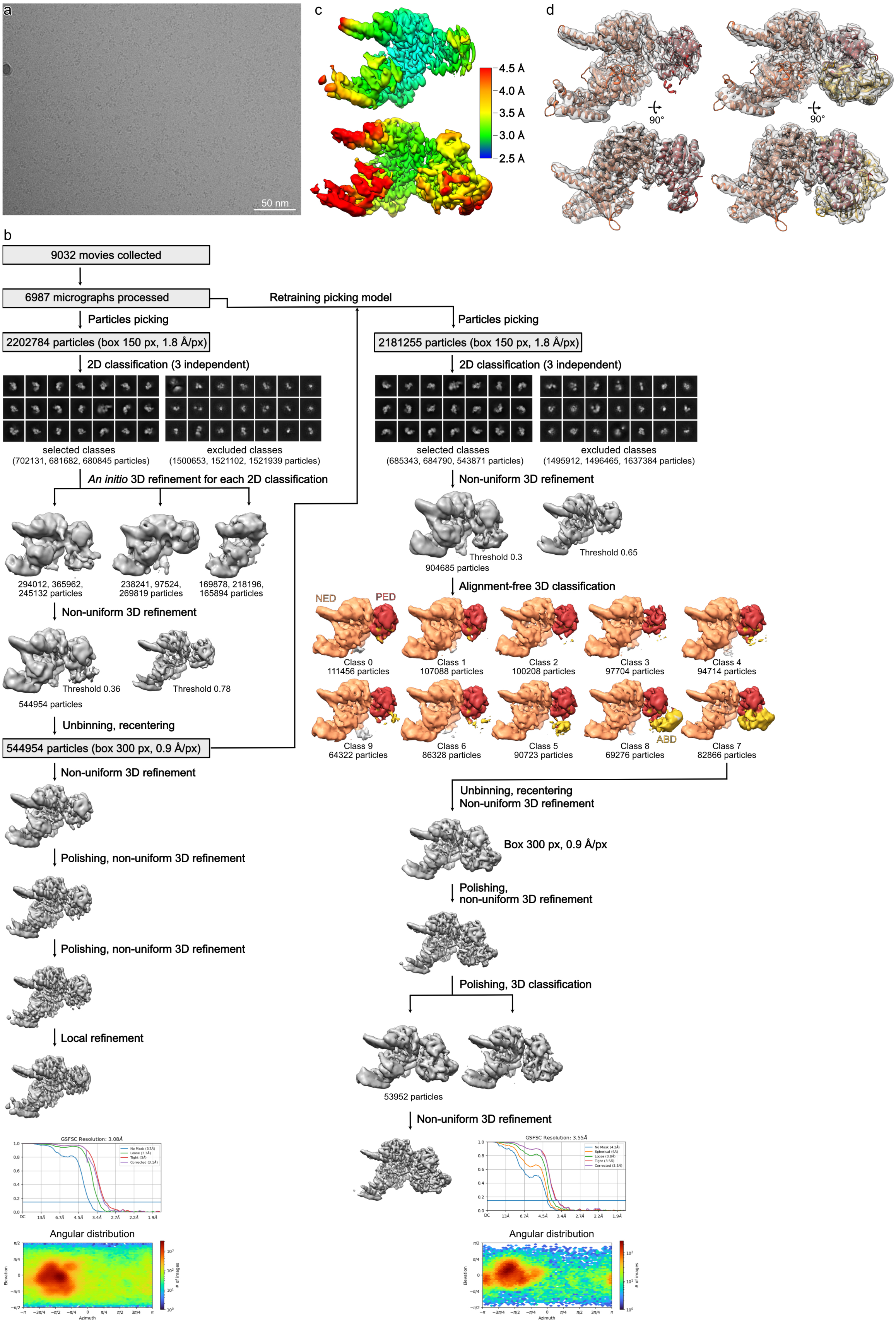

**Supplementary Fig. 7.** Processing of the Mcf1$_{C1397A\ \Delta15C}$ dataset. **a** An example of analyzed cryo-EM micrographs. **b** Processing overview with angular distributions and Fourier shell correlation curves of the final reconstructions used for model building. **c** Local resolution gradient of the final reconstructions. **d** Fit of the molecular models into the cryo-EM densities.

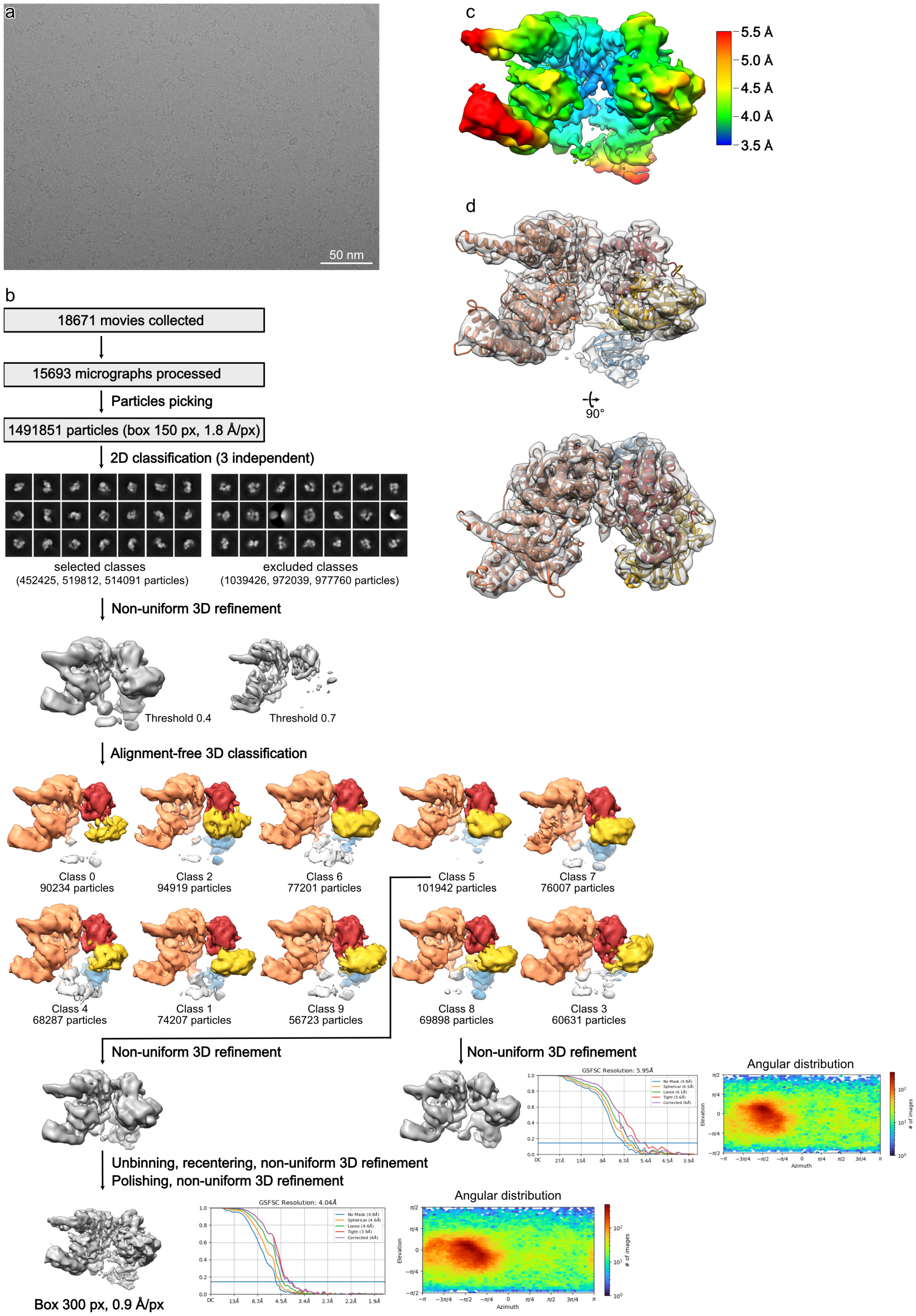

**Supplementary Fig. 8.** Processing of the Mcf1$_{C1397A\ \Delta15C}$-Arf3 complex dataset. **a** An example of analyzed cryo-EM micrographs. **b** Processing overview with angular distribution and Fourier shell correlation curves of the final reconstruction used for model building. **c** Local resolution gradient of the final reconstruction. **d** Fit of the molecular model into the cryo-EM density.

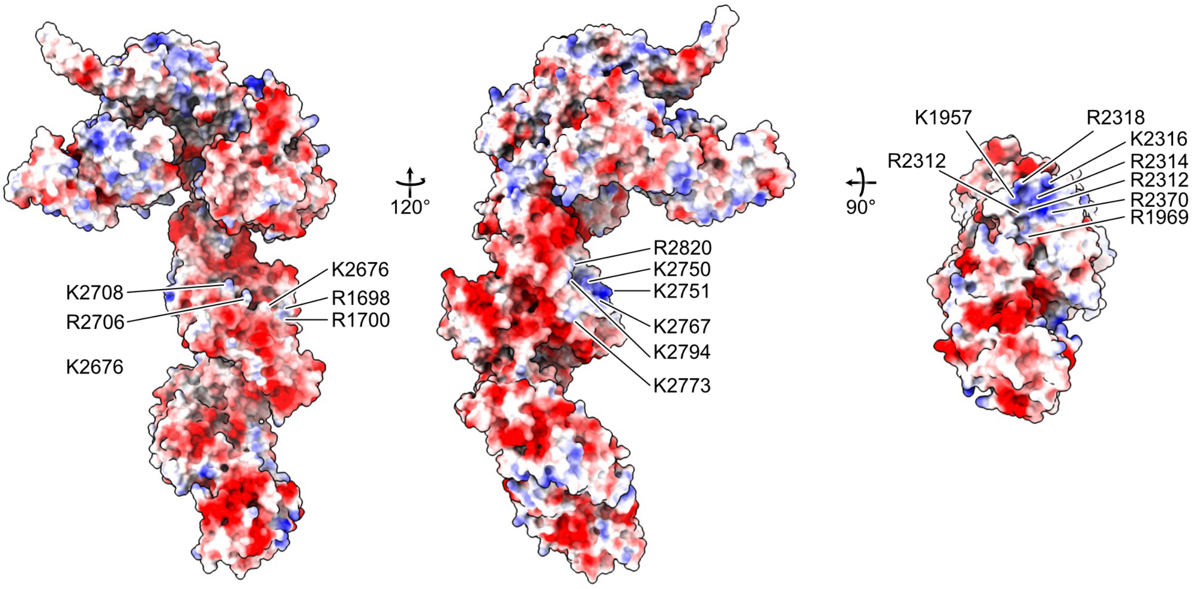

**Supplementary Fig. 9.** Surface of Mcf1 coloured by electrostatic Coulomb potential from $-10$ kcal mol$^{-1}$ (red) to +10 kcal mol$^{-1}$ (blue).

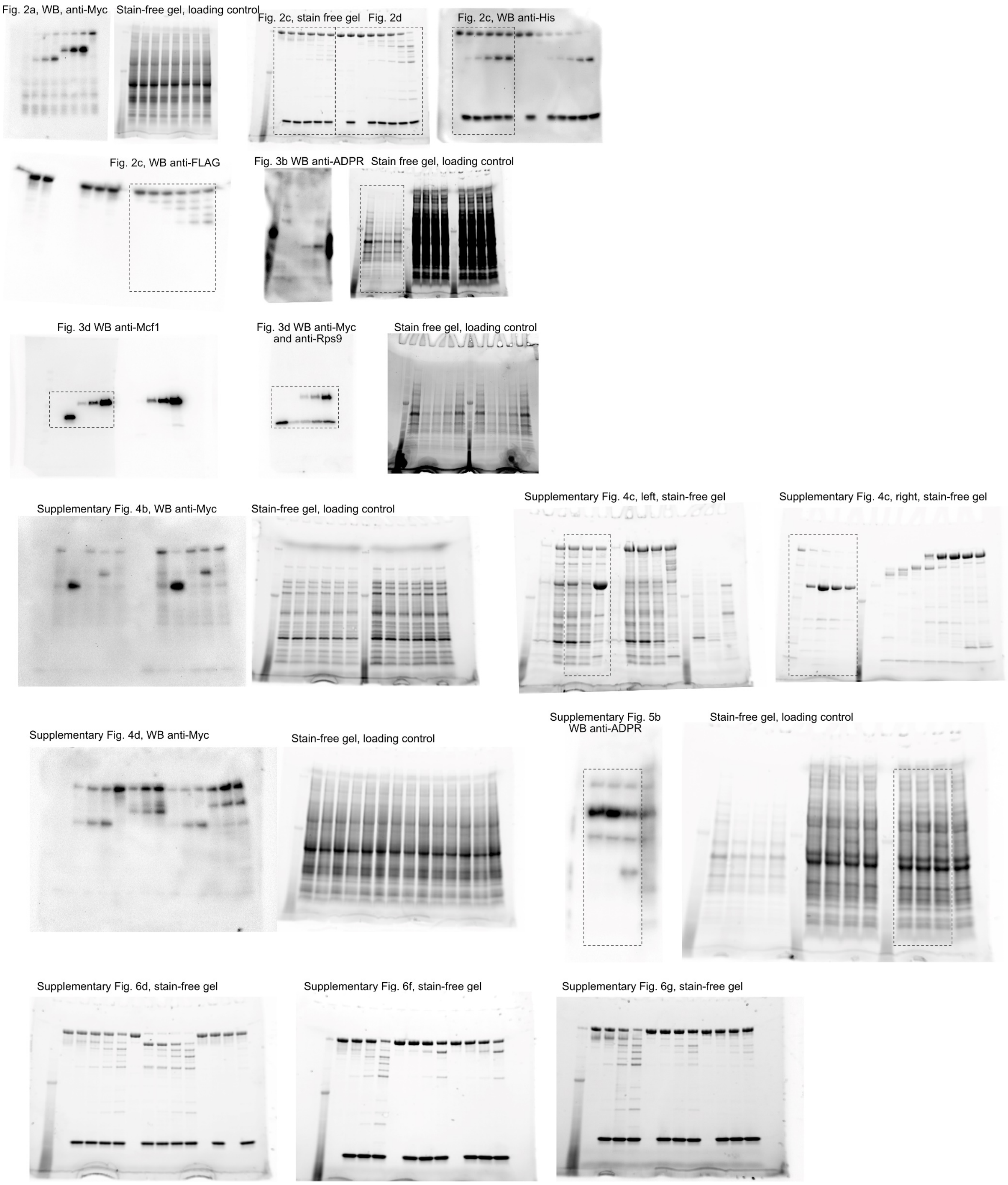

**Supplementary Fig. 10.** Uncropped SDS-PAGE, western blots (WB) and their loading controls.

**Supplementary Table 1. Cryo-EM data collection, refinement, and validation statistics**

| Sample | Full-length Mcf1 | Mcf1$_{C1397A\ \Delta15C}$ | Mcf1$_{C1397A\ \Delta15C}$-Arf3 complex |
|---|---|---|---|
| Microscope | Titan Krios | | |
| Voltage (kV) | 300 | | |
| Defocus range (μm) | -1 to -2.5 | | |
| Camera | Gatan K3 (super resolution mode) | | |
| Pixel size (Å) | 0.45 super resolution mode; 0.9 physical pixel size | | |
| Total electron dose (e/Å$^2$) | 71 and 69 | 60.85 | 62.6 |
| Exposure time, s | 3 | 3.5 | 2 |
| Frames per movie | 60 | 60 | 60 |
| Number of movies | 21,741 | 9,032 | 18,671 |
| **3D Refinement** | | | |
| Number of particles | 363,044 for building the tail region; 219,318 for building the head domain | 53,952 | 101,942 |
| Final resolution (Å) | 3.6 | 3.5 | 4.0 |
| **Atomic model statistics** | | | |
| Non-hydrogen atoms | 22,783 | 12,619 | 13,714 |
| Molprobity score | 1.52 | 1.51 | 1.68 |
| Clashscore | 3.44 | 4.21 | 6.41 |
| Cβ deviations >0.25Å | 0 | 0 | 0 |
| Bad bonds | 0 | 0 | 0 |
| Bad angles | 0 | 0 | 0 |
| Poor rotamers (%) | 0 | 0 | 0 |
| Favored rotamers (%) | 99.71 | 99.47 | 99.66 |
| Ramachandran favored (%) | 94.40 | 95.61 | 95.32 |
| Ramachandran allowed (%) | 5.57 | 4.39 | 4.68 |
| Ramachandran outliers (%) | 0.03 | 0 | 0 |

**Supplementary Table 2**. **List of primers, strains and plasmids used in this study.**

| Bacterial and yeast strains | Description | Reference |
|---|---|---|
| *E. coli* DH5α | $F^-$ Φ80*lac*ZΔM15 Δ(*lac*ZYA-*arg*F) U169 *rec*A1 *end*A1 *hsd*R17($r_k^-$, $m_k^+$) *pho*A *sup*E44 *thi*-1 *gyr*A96 *rel*A1 $\lambda^-$ | Invitrogen |
| *E. coli* BL21 (DE3) CodonPlus RIPL | $F^-$ *ompT hsdS*($r_B^-$ $m_B^-$) *dcm*$^+$ Tet$^r$ *gal*λ(DE3) *endA* Hte [*argU proL* Cam$^r$] [*argU ileY leuW* Strep/Spec$^r$] | Agilent |
| *S. cerevisiae* MH272-3fα | "Wild-type" strain, *ura3, leu2, his3, trp1, ade2* | [10] |
| *S. cerevisiae* Y388 | *S. cerevisiae* MH272-3fα + Iota-A [Ade] (p1323) | [11] |
| *S. cerevisiae* Y395 | *S. cerevisiae* MH272-3fα + empty vector [Ade] (2473) | [12] |
| *S. cerevisiae* Y401 | *S. cerevisiae* MH272-3fα + Mcf1 fragment 6 (amino acids 1133-1619) [Ade] (pB380) | This study |
| *S. cerevisiae* Y438 | *S. cerevisiae* MH272-3fα + Mcf1 fragment 6 D1505A [Ade] (pB408) | This study |
| *S. cerevisiae* Y439 | *S. cerevisiae* MH272-3fα + Mcf1 fragment 6 C1397A [Ade] (pB409) | This study |
| *S. cerevisiae* Y452 | *S. cerevisiae* MH272-3fα + Mcf1 fragment 6 H1486A [Ade] (pB419) | This study |
| *S. cerevisiae* Y639 | *S. cerevisiae* MH272-3fα + TcHVR ΔN100 (ART) [Ade] (pB631) | [13] |
| *S. cerevisiae* Y772 | *S. cerevisiae* MH272-3fα + Mcf1 fragment (amino acids 1-881) [Ade] (pB798) | This study |
| *S. cerevisiae* Y773 | *S. cerevisiae* MH272-3fα + Mcf1 fragment (amino acids 1-431) [Ade] (pB799) | This study |
| *S. cerevisiae* Y774 | *S. cerevisiae* MH272-3fα + Mcf1 fragment (amino acids 432-881) [Ade] (pB800) | This study |
| *S. cerevisiae* Y775 | *S. cerevisiae* MH272-3fα + Mcf1 fragment (amino acids 216-431) [Ade] (pB801) | This study |
| *S. cerevisiae* Y776 | *S. cerevisiae* MH272-3fα + Mcf1 fragment (amino acids 1-215) [Ade] (pB802) | This study |
| *S. cerevisiae* Y788 | *S. cerevisiae* MH272-3fα + Mcf1 fragment (amino acids 1-980) [Ade] (pB827) | This study |
| *S. cerevisiae* Y789 | *S. cerevisiae* MH272-3fα + Mcf1 fragment (amino acids 1-1080) [Ade] (pB828) | This study |
| *S. cerevisiae* Y790 | *S. cerevisiae* MH272-3fα + Mcf1 fragment (amino acids 1-1180) [Ade] (pB829) | This study |
| *S. cerevisiae* Y791 | *S. cerevisiae* MH272-3fα + Mcf1 fragment (amino acids 1-1280) [Ade] (pB830) | This study |
| *S. cerevisiae* Y792 | *S. cerevisiae* MH272-3fα + Mcf1 fragment (amino acids 1-1380) [Ade] (pB831) | This study |
| *S. cerevisiae* Y793 | *S. cerevisiae* MH272-3fα + Mcf1 fragment (amino acids 1-1480) [Ade] (pB832) | This study |
| *S. cerevisiae* Y794 | *S. cerevisiae* MH272-3fα + Mcf1 fragment (amino acids 1-1580) [Ade] (pB833) | This study |
| *S. cerevisiae* Y795 | *S. cerevisiae* MH272-3fα + Mcf1 fragment (amino acids 1-1630) [Ade] (pB834) | This study |
| *S. cerevisiae* Y836 | *S. cerevisiae* MH272-3fα + *Burkholderia* lethal factor 1 [Ade] (pB990) | This study |
| *S. cerevisiae* H009 | *S. cerevisiae* MH272-3fα + Mcf1 effector (amino acids 32-881) [Ade] (pH009) | This study |
| *S. cerevisiae* H010 | *S. cerevisiae* MH272-3fα + Mcf1 effector (amino acids 66-881) [Ade] (pH010) | This study |
| *S. cerevisiae* H011 | *S. cerevisiae* MH272-3fα + Mcf1 effector (amino acids 92-881) [Ade] (pH011) | This study |
| *S. cerevisiae* H012 | *S. cerevisiae* MH272-3fα + Mcf1 effector (amino acids 140-881) [Ade] (pH012) | This study |
| *S. cerevisiae* H013 | *S. cerevisiae* MH272-3fα + Mcf1 effector (amino acids 176-881) [Ade] (pH013) | This study |
| *S. cerevisiae* H014 | *S. cerevisiae* MH272-3fα + Mcf1 effector (amino acids 1-834) [Ade] (pH014) | This study |
| *S. cerevisiae* H016 | *S. cerevisiae* MH272-3fα + Mcf1 effector (amino acids 1-764) [Ade] (pH016) | This study |
| *S. cerevisiae* H017 | *S. cerevisiae* MH272-3fα + Mcf1 effector (amino acids 1-735) [Ade] (pH017) | This study |
| *S. cerevisiae* H018 | *S. cerevisiae* MH272-3fα + Mcf1 effector (amino acids 1-699) [Ade] (pH018) | This study |
| *S. cerevisiae* H023 | *S. cerevisiae* MH272-3fα + ABD (amino acids 913-1125) [Ade] (pH023) | This study |
| *S. cerevisiae* H024 | *S. cerevisiae* MH272-3fα + ABD G1031C (amino acids 913-1125) [Ade] (pH024) | This study |

| **Plasmids for experiments in *S. cerevisiae*** | | |
|---|---|---|
| 2473 YEpGal555 | *E. coli*/*S. cerevisiae* shuttle vector [ADE2] with Gal1 promoter | [12] |
| pB380 YEpGal555 Mcf1 fragment 6 | Fragment of the *mcf1* gene, encoding amino acids 1133-1619, was PCR amplified using oligonucleotides TATACTCGAGCAATTCGTCGCCTACCGCGAGC and TATATAGCTAGCGAACCGTGCCCCCCATTGCTCG from pB394, digested with XhoI and NheI and ligated in the digested 2473 YEpGal555 vector. | This study |
| pB408 YEpGal555 Mcf1 fragment 6 D1505A | The mutation D1505A was introduced by two-step overlap PCR using oligonucleotides TATACTCGAGCAATTCGTCGCCTACCGCGAGC, TATATAGCTAGCGAACCGTGCCCCCCATTGCTCG, CTACTTTTATGCCCCGAACGTCG, and CGACGTTCGGGGCATAAAAGTAG and pB380 as the template. The PCR product was digested with XhoI and NheI, and ligated into 2473 YEpGal555 vector. | This study |
| pB409 YEpGal555 Mcf1 fragment 6 C1397A | The mutation C1397A was introduced by two-step overlap PCR using oligonucleotides TATACTCGAGCAATTCGTCGCCTACCGCGAGC, TATATAGCTAGCGAACCGTGCCCCCCATTGCTCG, GTGACCGATCTGGTGGCCGTGCTTATCCTTTGGTCAGGGCC, and CTGACCAAAGGATAAGCACGGCCACCAGATCGGTCACCGAC and pB380 as the template. The PCR product was digested with XhoI and NheI, and ligated into 2473 YEpGal555 vector. | This study |
| pB419 YEpGal555 Mcf1 fragment 6 H1486A | The mutation H1486A was introduced by two-step overlap PCR using oligonucleotides TATACTCGAGCAATTCGTCGCCTACCGCGAGC, TATATAGCTAGCGAACCGTGCCCCCCATTGCTCG, CAATACCCAGAATGCCTCGATGATGG, and CCATCATCGAGGCATTCTGGGTATTG and pB380 as the template. The PCR product was digested with XhoI and NheI, and ligated into 2473 YEpGal555 vector. | This study |
| pB631 YEpGal555 TcART | TccC3 with a 100 amino-acid long N-terminal deletion in 2473 YEpGal555 vector. | [13] |
| pB798 YEpGal555 Mcf1 1-881 | Fragment of the *mcf1* gene, encoding amino acids 1-881, was PCR amplified using oligonucleotides TATAGGATCCATGGAACAGAAGTTGATTTCCGAAGAAGACCTCGCTTCTATATCCAAAGATTTTACG and ATATCTCGAGGCCCTGATTGATGATCGGCAAG from pB394, digested with BamHI and XhoI and ligated in the digested 2473 YEpGal555 vector. | This study |
| pB799 YEpGal555 Mcf1 1-431 | Fragment of the *mcf1* gene, encoding amino acids 1-431, was PCR amplified using oligonucleotides TATAGGATCCATGGAACAGAAGTTGATTTCCGAAGAAGACCTCGCTTCTATATCCAAAGATTTTACG and ATATCTCGAGACGACCGGCAAAACCTAC from pB394, digested with BamHI and XhoI and ligated in the digested 2473 YEpGal555 vector. | This study |
| pB800 YEpGal555 Mcf1 432-881 | Fragment of the *mcf1* gene, encoding amino acids 432-881, was PCR amplified using oligonucleotides tatagGATCCatggaacagaagttgatttccgaagaagacctcCAGTACTTGCTGGAAATGCC and ATATCTCGAGGCCCTGATTGATGATCGGCAAG from pB394, digested with BamHI and XhoI and ligated in the digested 2473 YEpGal555 vector. | This study |
| pB801 YEpGal555 Mcf1 216-431 | Fragment of the *mcf1* gene, encoding amino acids 216-431, was PCR amplified using oligonucleotides tatagGATCCatggaacagaagttgatttccgaagaagacctcAAGGTGCCTCAGGCCTCCG and ATATCTCGAGACGACCGGCAAAACCTAC from pB394, digested with BamHI and XhoI and ligated in the digested 2473 YEpGal555 vector. | This study |
| pB802 YEpGal555 Mcf1 1-215 | Fragment of the *mcf1* gene, encoding amino acids 1-215, was PCR amplified using oligonucleotides | This study |

| | | |
|---|---|---|
| | TATAGGATCCATGGAACAGAAGTTGATTTCCGAAGAAGACCTCGCTTCTATATCCAAAGATTTTACG and ATATCTCGAGGGTCAGCGCACCGAGAGC from pB394, digested with BamHI and XhoI and ligated in the digested 2473 YEpGal555 vector. | |
| pB827 YEpGal555 Mcf1 1-980 | Fragment of the *mcf1* gene, encoding amino acids 1-980, was PCR amplified using oligonucleotides TATAGGATCCATGGAACAGAAGTTGATTTCCGAAGAAGACCTCGCTTCTATATCCAAAGATTTTACG and tatactcgaGCTCAGTGTACTCAAGCTCATACAAG from pB394, digested with BamHI and XhoI and ligated in the digested 2473 YEpGal555 vector. | This study |
| pB828 YEpGal555 Mcf1 1-1080 | Fragment of the *mcf1* gene, encoding amino acids 1-1080, was PCR amplified using oligonucleotides TATAGGATCCATGGAACAGAAGTTGATTTCCGAAGAAGACCTCGCTTCTATATCCAAAGATTTTACG and tatactcgaGCGCAGCGAGGCCCTCCGCGG from pB394, digested with BamHI and XhoI and ligated in the digested 2473 YEpGal555 vector. | This study |
| pB829 YEpGal555 Mcf1 1-1180 | Fragment of the *mcf1* gene, encoding amino acids 1-1180, was PCR amplified using oligonucleotides TATAGGATCCATGGAACAGAAGTTGATTTCCGAAGAAGACCTCGCTTCTATATCCAAAGATTTTACG and tatactcgaGGGCCCACGCAGCAATCGCCGG from pB394, digested with BamHI and XhoI and ligated in the digested 2473 YEpGal555 vector. | This study |
| pB830 YEpGal555 Mcf1 1-1280 | Fragment of the *mcf1* gene, encoding amino acids 1-1280, was PCR amplified using oligonucleotides TATAGGATCCATGGAACAGAAGTTGATTTCCGAAGAAGACCTCGCTTCTATATCCAAAGATTTTACG and tatactcgaGATCGGTGCGCTGCTTACCCC from pB394, digested with BamHI and XhoI and ligated in the digested 2473 YEpGal555 vector. | This study |
| pB831 YEpGal555 Mcf1 1-1380 | Fragment of the *mcf1* gene, encoding amino acids 1-1380, was PCR amplified using oligonucleotides TATAGGATCCATGGAACAGAAGTTGATTTCCGAAGAAGACCTCGCTTCTATATCCAAAGATTTTACG and tatactcgaGTACCAAGCGGTCAAAGACACTGCC from pB394, digested with BamHI and XhoI and ligated in the digested 2473 YEpGal555 vector. | This study |
| pB832 YEpGal555 Mcf1 1-1480 | Fragment of the *mcf1* gene, encoding amino acids 1-1480, was PCR amplified using oligonucleotides TATAGGATCCATGGAACAGAAGTTGATTTCCGAAGAAGACCTCGCTTCTATATCCAAAGATTTTACG and tatactcgaGCGCGAACATCGAGGTTCCGG from pB394, digested with BamHI and XhoI and ligated in the digested 2473 YEpGal555 vector. | This study |
| pB833 YEpGal555 Mcf1 1-1580 | Fragment of the *mcf1* gene, encoding amino acids 1-1580, was PCR amplified using oligonucleotides TATAGGATCCATGGAACAGAAGTTGATTTCCGAAGAAGACCTCGCTTCTATATCCAAAGATTTTACG and tatactcgaGGACGCTGGACAGTTCCTCAAATC from pB394, digested with BamHI and XhoI and ligated in the digested 2473 YEpGal555 vector. | This study |
| pB834 YEpGal555 Mcf1 1-1630 | Fragment of the *mcf1* gene, encoding amino acids 1-1630, was PCR amplified using oligonucleotides TATAGGATCCATGGAACAGAAGTTGATTTCCGAAGAAGACCTCGCTTCTATATCCAAAGATTTTACG and tatactcgaGGTGCTCTTGCGCCAGCCGGG from pB394, digested with BamHI and XhoI and ligated in the digested 2473 YEpGal555 vector. | This study |
| pB990 YEpGal555 *Burkholderia* lethal factor 1 | Gene, encoding *Burkholderia pseudomallei* lethal factor 1 (Blf1, Uniprot Q63UP7), was codon-optimized for expression in *S. cerevisiae*, synthesized and cloned into 2473 YEpGal555 vector by XhoI and KpnI sites. | This study |
| pH009 YEpGal555 Mcf1 32-881 | Fragment of the *mcf1* gene, encoding amino acids 32-881, was PCR amplified using oligonucleotides tatagGATCCatggaacagaagttgatttccgaagaagacctcATGAGTCCTGATGAGCGTAC | This study |

| | | |
|---|---|---|
| | and ATATCTCGAGGCCCTGATTGATGATCGGCAAG from pB394, digested with BamHI and XhoI and ligated in the digested 2473 YEpGal555 vector. | |
| pH010 YEpGal555 Mcf1 66-881 | Fragment of the *mcf1* gene, encoding amino acids 66-881, was PCR amplified using oligonucleotides tatagGATCCatggaacagaagttgatttccgaagaagacctcATGCAGGACAACCCC and ATATCTCGAGGCCCTGATTGATGATCGGCAAG from pB394, digested with BamHI and XhoI and ligated in the digested 2473 YEpGal555 vector. | This study |
| pH011 YEpGal555 Mcf1 92-881 | Fragment of the *mcf1* gene, encoding amino acids 92-881, was PCR amplified using oligonucleotides tatagGATCCatggaacagaagttgatttccgaagaagacctcGGCACGCCGGC and ATATCTCGAGGCCCTGATTGATGATCGGCAAGfrom pB394, digested with BamHI and XhoI and ligated in the digested 2473 YEpGal555 vector. | This study |
| pH012 YEpGal555 Mcf1 140-881 | Fragment of the *mcf1* gene, encoding amino acids 140-881, was PCR amplified using oligonucleotides tatagGATCCatggaacagaagttgatttccgaagaagacctcAATGCGACGACATTGAGAG and ATATCTCGAGGCCCTGATTGATGATCGGCAAG from pB394, digested with BamHI and XhoI and ligated in the digested 2473 YEpGal555 vector. | This study |
| pH013 YEpGal555 Mcf1 176-881 | Fragment of the *mcf1* gene, encoding amino acids 176-881, was PCR amplified using oligonucleotides tatagGATCCatggaacagaagttgatttccgaagaagacctcAGCCAAAAATATGTGACGC and ATATCTCGAGGCCCTGATTGATGATCGGCAAG from pB394, digested with BamHI and XhoI and ligated in the digested 2473 YEpGal555 vector. | This study |
| pH014 YEpGal555 Mcf1 1-834 | Fragment of the *mcf1* gene, encoding amino acids 1-834, was PCR amplified using oligonucleotides tatagGATCCatggaacagaagttgatttccgaagaagacctcGCTTCTATATCCAAAGATTTTAC G and atatCTCGAGttaTTCATAACGCGCAGCG from pB394, digested with BamHI and XhoI and ligated in the digested 2473 YEpGal555 vector. | This study |
| pH016 YEpGal555 Mcf1 1-764 | Fragment of the *mcf1* gene, encoding amino acids 1-764, was PCR amplified using oligonucleotides tatagGATCCatggaacagaagttgatttccgaagaagacctcGCTTCTATATCCAAAGATTTTAC G and atatCTCGAGttaGTCATTCTTGAAGTCCGG from pB394, digested with BamHI and XhoI and ligated in the digested 2473 YEpGal555 vector. | This study |
| pH017 YEpGal555 Mcf1 1-735 | Fragment of the *mcf1* gene, encoding amino acids 1-735, was PCR amplified using oligonucleotides tatagGATCCatggaacagaagttgatttccgaagaagacctcGCTTCTATATCCAAAGATTTTAC G and atatCTCGAGttaCTCAGAGCTGGTATTGAG from pB394, digested with BamHI and XhoI and ligated in the digested 2473 YEpGal555 vector. | This study |
| pH018 YEpGal555 Mcf1 1-699 | Fragment of the *mcf1* gene, encoding amino acids 1-699, was PCR amplified using oligonucleotides tatagGATCCatggaacagaagttgatttccgaagaagacctcGCTTCTATATCCAAAGATTTTAC G and atatCTCGAGttaCTTGGAAAAGCTATCGGC from pB394, digested with BamHI and XhoI and ligated in the digested 2473 YEpGal555 vector. | This study |
| pH023 YEpGal555 Mcf1 913-1125 | Fragment of the *mcf1* gene, encoding amino acids 913-1125, was PCR amplified using oligonucleotides tatagGATCCatggaacagaagttgatttccgaagaagacctcGCCGGGTTGACGTCTGTAGG and atatCTCGAGttaTTGACTGTCCGCCACCTGGATC from pB397, digested with BamHI and XhoI and ligated in the digested 2473 YEpGal555 vector. | This study |

| | | |
|---|---|---|
| pH024 YEpGal555 Mcf1 913-1125 G1031C | The mutation G1031C was introduced by two-step overlap PCR using oligonucleotides CGCTATCGGGCTGCTCG, CGAGCAGCCCGATAGCG, tatagGATCCatggaacagaagttgatttccgaagaagacctcGCCGGGTTGACGTCTGTAGG and atatCTCGAGttaTTGACTGTCCGCCACCTGGATC and pB397 as the template. The PCR product was digested with BamHI and XhoI and ligated into digested 2473 YEpGal555 | This study |
| **Plasmids for protein expression in *E. coli*** | | |
| pETDuet | *E. coli* vector for expression of two genes. | Novagen |
| 1330 pET28 Iota-A | *Clostridium perfringens* Iota-A with N-terminal His-tag in pET28 vector | [11] |
| pB656 pET28 MBP-TcHVR ΔN100 (TcART) | TcHVR with the N-terminal deletion in fusion with MBP tag in pET28 vector | [13] |
| pB475 pET28 His-Arf3 Q71L | Gene, encoding human ADP-ribosylating factor 3 (Arf3; Uniprot P61204) with N-terminal deletion of 17 amino acids and mutation Q71L, was codon-optimized for expression *E. coli*, synthesized and cloned into pET28a vector by NdeI and BamHI sites. | This study |
| pB394 pET28 His-Mcf1 | *Mcf1* gene (Uniprot Q8KT65) was PCR amplified from *Photorhabdus luminescens* strain akhurstii using oligonucleotides tatacatatgGCTTCTATATCCAAAGATTTTACGAAC and tatagagctcTTAGATGGCCCAAGGCAGCTCGA. The PCR product was digested with NdeI and SacI and inserted into digested pET28a vector. | This study |
| pB397 pET28 His-Myc-Mcf1-FLAG | N-terminal Myc and C-terminal FLAG tags were added to pB394 in three steps. First, we removed a stop codon at the end of *mcf1* gene in pB394 by PCR using oligonucleotides tatacatatgGCTTCTATATCCAAAGATTTTACGAAC and tatagagctcGATGGCCCAAGGCAGCTCGA and pB394 as the template. The PCR product was digested with NdeI and SacI and inserted into pET28a (resulting plasmid pB395).<br>Second, we annealed oligonucleotides encoding Myc-tag tagtgaacagaagttgatttccgaagaagacctctc and tagagaggtcttcttcggaaatcaacttctgttcac, and inserted them into NdeI-digested pB395 (resulting plasmid pB396)<br>Finally, we annealed oligonucleotides encoding FLAG-tag agattacaaggatgacgacgataagtaaagct and ttacttatcgtcgtcatccttgtaatctagct, and inserted into SacI-digested pB396. | This study |
| pB446 pET28 His-Myc-Mcf1_C1397A-FLAG | The mutation C1397A was introduced by two-step overlap PCR using oligonucleotides aactttaagaaggagatataccatgggcagcagc, CCGCAAGCTtgtcgacggagctttacttatcgtcgtc, GTGACCGATCTGGTGGCCGTGCTTATCCTTTGGTCAGGGCC, and CTGACCAAAGGATAAGCACGGCCACCAGATCGGTCACCGAC and pB397 as the template. The PCR product was digested with NcoI and HindIII, and ligated into digested pET28 vector. | This study |
| pB852 pETDuet Arf3 Q71L | Arf3 Q71L was PCR-amplified from pB475 with oligonucleotides TATACAtatgtcatggtctcatccacaattcgaaaagtcacgcattctgatggtgggcctggatgcgg and GCTAGTTATTGCTCAGCGG. PCR product was digested with NdeI and SalI, and ligated into NdeI/XhoI-digested pETDuet vector. | This study |

| | | |
|---|---|---|
| pB858 pETDuet Arf3 Q71L His-Myc-Mcf1-FLAG | *Mcf1* gene was cut by NcoI and HindIII from pB397 and inserted into digested pB852. | This study |
| pB891 pET28 His-Mcf1-Myc | C-terminal Myc-tag was added to Mcf1 by annealing oligonucleotides cggatcaggtgaacagaagttgatttccgaagaagacctcta and agcttagaggtcttcttcggaaatcaacttctgttcacctgatccgagct and inserting them into SacI/HindIII-digested pB395 | This study |
| pB914 His-Mcf1 Δ15C | C-terminal deletion was performed by PCR using oligonucleotides tatacatatgGCTTCTATATCCAAAGATTTTACGAAC and tatagagctcTTACCTGGTAAAACCCTTGACCAAGTG, and pB394 as the template. The PCR product was digested with NdeI and SacI and inserted into digested pET28a vector. | This study |
| pB915 His-Mcf1 C1397A Δ15C | The mutation C1397A was introduced by two-step overlap PCR using oligonucleotides tatacatatgGCTTCTATATCCAAAGATTTTACGAAC, tatagagctcTTACCTGGTAAAACCCTTGACCAAGTG, GTGACCGATCTGGTGGCCGTGCTTATCCTTTGGTCAGGGCC, and CTGACCAAAGGATAAGCACGGCCACCAGATCGGTCACCGAC and pB914 as the template. The PCR product was digested with NdeI and SacI and inserted into digested pET28a vector. | This study |
| pB916 pETDuet His-Myc-Mcf1-FLAG | *Mcf1* gene was cut by NcoI and HindIII from pB397 and inserted into the digested pETDuet vector. | This study |
| pB917 pETDuet Arf3 Q71L His-Myc-Mcf1_C1397A-FLAG | *Mcf1* gene was cut by NcoI and HindIII from pB446 and inserted into digested pB852. | This study |
| pB973 pET28 His-Mcf1 Y1286S Δ15C | The Y1286S mutation in the *mcf1* gene was introduced by two-step overlap PCR using oligonucleotides tatacatatgGCTTCTATATCCAAAGATTTTACGAAC, tatagagctcTTACCTGGTAAAACCCTTGACCAAGTG, GCGTGGGTGAACGTagCGCTATCTCTGAAG and CTTCAGAGATAGCGctACGTTCACCCACGC and pB914 as the template. The PCR product was digested with NdeI and SacI and inserted into digested pET28a vector. | This study |
| pB987 pET28 His-Mcf1 R957A Δ15C | The R957A mutation in the *mcf1* gene was introduced by two-step overlap PCR using oligonucleotides tatacatatgGCTTCTATATCCAAAGATTTTACGAAC, tatagagctcTTACCTGGTAAAACCCTTGACCAAGTG, CTGGTCAGATCGGGGgcCCTCCCGGCAGAGGG and CCCTCTGCCGGGAGGgcCCCCGATCTGACCAG and pB914 as the template. The PCR product was digested with NdeI and SacI and inserted into digested pET28a vector. | This study |
| pB989 pET28 His-Mcf1 Y1286S C1397A Δ15C | The C1397A mutation in the Mcf1 Y1286S variant was introduced by two-step overlap PCR using oligonucleotides tatacatatgGCTTCTATATCCAAAGATTTTACGAAC, tatagagctcTTACCTGGTAAAACCCTTGACCAAGTG, GTGACCGATCTGGTGGCCGTGCTTATCCTTTGGTCAGGGCC and CTGACCAAAGGATAAGCACGGCCACCAGATCGGTCACCGAC and pB973 as the template. The PCR product was digested with NdeI and SacI and inserted into digested pET28a vector. | This study |
| pB992 pET28 His-Mcf1 L1359G Δ15C | The L1359G mutation in the *mcf1* gene was introduced by two-step overlap PCR using oligonucleotides tatacatatgGCTTCTATATCCAAAGATTTTACGAAC, tatagagctcTTACCTGGTAAAACCCTTGACCAAGTG, CTGAGTACATCGATAAGGTAggGCGGCAGACTGCGG and | This study |

| | | |
|---|---|---|
| | CCGCAGTCTGCCGCccTACCTTATCGATGTACTCAG and pB914 as the template. The PCR product was digested with NdeI and SacI and inserted into digested pET28a vector. | |
| pB993 pET28 His-Mcf1 L1359G R1360G Δ15C | The L1359G and R1360G mutations in the *mcf1* gene were introduced by two-step overlap PCR using oligonucleotides tatacatatgGCTTCTATATCCAAAGATTTTACGAAC, tatagagctcTTACCTGGTAAAACCCTTGACCAAGTG, CTGAGTACATCGATAAGGTAggGgGGCAGACTGCGGTATTC and GAATACCGCAGTCTGCCcCccTACCTTATCGATGTACTCAG and pB914 as the template. The PCR product was digested with NdeI and SacI and inserted into digested pET28a vector. | This study |
| pB996 pET28 His-Mcf1 L911A K912A Δ15C | The L911A K912A mutations in the *mcf1* gene were introduced by two-step overlap PCR using oligonucleotides tatacatatgGCTTCTATATCCAAAGATTTTACGAAC, tatagagctcTTACCTGGTAAAACCCTTGACCAAGTG, GGCGGTGAAGGATCTCGAAgccgcgGCCGGGTTGACGTCTGTAGG and CCTACAGACGTCAACCCGGCcgcggcTTCGAGATCCTTCACCGCC and pB914 as the template. The PCR product was digested with NdeI and SacI and inserted into digested pET28a vector. | This study |
| pB1000 pETDuet His-Myc-Mcf1-FLAG L911A K912A | The mutations L911A K912A were introduced by two-step overlap PCR using oligonucleotides aactttaagaaggagatataccatgggcagcagc, CCGCAAGCTtgtcgacggagctttacttatcgtcgtc, GGCGGTGAAGGATCTCGAAgccgcgGCCGGGTTGACGTCTGTAGG, and CCTACAGACGTCAACCCGGCcgcggcTTCGAGATCCTTCACCGCC and pB397 as the template. The PCR product was digested with NcoI and HindIII, and ligated into digested pETDuet vector. | This study |
| pB1004 pET28 His-Myc-Mcf1-FLAG L911A K912A I1271A Q1272A G1273A | The mutations I1271A Q1272A G1273A were introduced by two-step overlap PCR using oligonucleotides aactttaagaaggagatataccatgggcagcagc, CCGCAAGCTtgtcgacggagctttacttatcgtcgtc, CGGATTGACGATACAGCCgccgcagcCGGGGGTAAGCAGCGC and GCGCTGCTTACCCCCGgctgcggcGGCTGTATCGTCAATCCG and pB1000 as the template. The PCR product was digested with NcoI and HindIII, and ligated into digested pET28 vector. | This study |
| pB1006 pETDuet Arf3 Q71L His-Myc-Mcf1-FLAG L911A K912A | *Mcf1* gene with required mutations was cut by NcoI and HindIII from pB1000 and inserted into the digested pB852 vector. | This study |
| pB1008 pETDuet Arf3 Q71L His-Myc-Mcf1-FLAG L911A K912A I1271A Q1272A G1273A | *Mcf1* gene with required mutations was cut by NcoI and HindIII from pB1004 and inserted into the digested pB852 vector. | This study |
| pB1012 pET28 His-Myc-Mcf1-FLAG I1271A Q1272A G1273A | The mutations I1271A Q1272A G1273A were introduced by two-step overlap PCR using oligonucleotides aactttaagaaggagatataccatgggcagcagc, CCGCAAGCTtgtcgacggagctttacttatcgtcgtc, CGGATTGACGATACAGCCgccgcagcCGGGGGTAAGCAGCGC and GCGCTGCTTACCCCCGgctgcggcGGCTGTATCGTCAATCCG and pB397 as the template. The PCR product was digested with NcoI and HindIII, and ligated into digested pET28 vector. | This study |

| | | |
|---|---|---|
| pB1022 pET28 His-Mcf1 ΔK907 Δ15C | The ΔK907 deletion in the *mcf1* gene was introduced by two-step overlap PCR using oligonucleotides tatacatatgGCTTCTATATCCAAAGATTTTACGAAC, tatagagctcTTACCTGGTAAAACCCTTGACCAAGTG, GCTGTTGGGGCAGGCGGTGGATCTCGAACTCAAG and CTTGAGTTCGAGATCCACCGCCTGCCCCAACAGC and pB914 as the template. The PCR product was digested with NdeI and SacI and inserted into digested pET28a vector. | This study |
| pB1023 pET28 His-Mcf1 ΔK907 ΔD908 Δ15C | The ΔK907 and ΔD908 deletions in the *mcf1* gene were introduced by two-step overlap PCR using oligonucleotides tatacatatgGCTTCTATATCCAAAGATTTTACGAAC, tatagagctcTTACCTGGTAAAACCCTTGACCAAGTG, CGCTGTTGGGGCAGGCGGTGCTCGAACTCAAGGCCG and CGGCCTTGAGTTCGAGCACCGCCTGCCCCAACAGCG and pB914 as the template. The PCR product was digested with NdeI and SacI and inserted into digested pET28a vector. | This study |
